# MG53 slows neuromuscular junction loss and prolongs survival in ALS

**DOI:** 10.1101/2021.04.22.441038

**Authors:** Jianxun Yi, Ang Li, Xuejun Li, Ki Ho Park, Xinyu Zhou, Frank Yi, Yajuan Xiao, Dosuk Yoon, Tao Tan, Lyle W. Ostrow, Jianjie Ma, Jingsong Zhou

## Abstract

Respiratory failure from progressive respiratory muscle weakness is the most common cause of death in amyotrophic lateral sclerosis (ALS). Defects in neuromuscular junctions (NMJs) and progressive NMJ loss occur at early stages, thus stabilizing and preserving NMJs represents a potential therapeutic strategy to slow ALS disease progression. Here we demonstrate that NMJ damage is repaired by MG53, an intrinsic muscle protein involved in plasma membrane repair. Compromised diaphragm muscle membrane repair and NMJ integrity are early pathological findings in ALS. Diaphragm muscles from ALS mouse models show increased susceptibility to injury and intracellular MG53 aggregation, which is also a hallmark of human muscle samples from ALS patients. We show that systemic administration of recombinant human MG53 protein (rhMG53) in ALS mice protects against injury to diaphragm muscle, preserves NMJ integrity, and slows ALS disease progression. As MG53 is present in circulation in rodents and humans under physiological conditions, our findings provide proof-of-concept data supporting MG53 as a potentially safe and effective therapy to mitigate ALS progression.

## Introduction

Amyotrophic lateral sclerosis (ALS) is a fatal neuromuscular disease characterized by progressive motor neuron loss and muscle atrophy[1, 2]. Progressive respiratory muscle weakness is a main cause of morbidity and eventual death[3, 4]. ALS appears to be a combination of “dying-forward” and/or “dying-back” pathophysiological processes, starting in cortical motor neurons and glia, or at the muscle and NMJ[5–16]. It is possible that the balance of these processes differs in subsets of ALS patients, and several mechanisms have been proposed for how they may interrelate. For example, NMJ degeneration is associated with mitochondrial dysfunction in ALS[17–21], which involves bidirectional crosstalk between myofibers and neuron[6, 8, 9, 17–21]. Therapeutic approaches that sustain NMJ integrity and muscle function could slow disease progression. Thus, it is critical to understand the molecular mechanisms associated with NMJ degeneration in ALS.

During respiration, the diaphragm constantly undergoes contraction-relaxation, a process that leads to injury to the muscle membrane. Inadequate repair of injury to the sarcolemma can disrupt NMJ integrity and contribute to diaphragm wasting in ALS. MG53, a member of the tripartite motif (TRIM) protein family[22], was identified as an essential component of the cell membrane repair machinery[23–26]. Genetic ablation of MG53 results in defective membrane repair and tissue regenerative capacity[23, 24, 27, 28]. A series of studies have shown that recombinant human MG53 (rhMG53) protein protects various cell types against membrane disruption when applied to the extracellular environment, and ameliorates pathology associated with muscular dystrophy[29], acute lung injury[30], myocardial infarction[31], acute kidney injury[32], and ischemic brain damage[33] in animal models.

In this study, we identify NMJ as an active site of injury-repair by MG53 and show that compromised diaphragm muscle membrane repair occurs prior to symptom onset in ALS mice. Oxidative stress associated with mitochondrial dysfunction in ALS muscle leads to formation of MG53 protein aggregates and disruption of MG53-mediated membrane repair. Diaphragm muscles from ALS mouse models show increased susceptibility to injury and intracellular MG53 aggregation, which is also seen in human muscle samples from ALS patients. Remarkably, systemic application of rhMG53 in ALS mice protects against injury to the diaphragm, preserves integrity of NMJ, and significantly alleviates ALS disease progression.

## Materials and Methods

### Animal models

The ALS transgenic mouse model (G93A) with genetic background of B6SJL was originally generated by Drs. Deng and Siddique’s group at Northwestern University[34], which was also deposited to the Jackson Lab as B6SJL-Tg (SOD1*G93A). We obtained this colony from Dr. Deng and maintained it through breeding the B6SJL/G93A male mice with B6SJL/WT female mice. All experiments were carried out in accordance with the recommendations in the Guide for the Care and Use of Laboratory Animals of the National Institutes of Health. Protocols on the usage of mice were approved by the Institutional Animal Care and Use Committee of University of Missouri at Kansas City, Kansas City University of Medicine and Biosciences and University of Texas at Arlington. Sprague-Dawley rats (4-month old) were purchased from Charles River Laboratories. Protocol on the usage of rats was approved by the Institutional Animal Care and Use Committee of the Ohio State University.

### Isolation of single live FDB muscle fibers (myofibers)

Experimental mice were euthanized by cervical dislocation, and FDB muscles were removed for enzyme digestion to obtain individual myofibers for functional or biochemical studies [18, 20, 21]. Briefly, FDB muscles were digested in modified Krebs solution (0 Ca^2+^) containing 0.2% type I Collagenase (Sigma), for 1 hour at 37 °C. After digestion, muscles were kept in collagenase-free Krebs solution (with 2.5 mM Ca^2+^ and 10 mM glucose) at 4°C and used for studies within 24 hours.

### Evaluation of T-tubule network integrity and mitochondrial membrane potential at NMJ

Freshly isolated FDB myofibers were seeded in laminin (Santa Cruz sc-29012) coated glass bottom dishes and stained with TMRE (1: 3000 dilution from 150 μM stock solution, Invitrogen T669), BTX-Alexa Fluor 488 (1: 1000 dilution from 1 mg/ml stock solution, Invitrogen B13422) and CellMask DeepRed (1:1000 dilution from 5 mg/ml stock solution, Invitrogen C10046) for 30 min. The myofibers were washed 5 times with 2.5 mM Ca^2+^Krebs solution before imaging with confocal microscopy. BTX-Alexa Fluor 488 was excited by 488 nm laser (emission filter 500-550 nm). TMRE was excited by 568 nm laser (emission filter 575-625 nm). CellMask DeepRed was excited by 633 nm laser (emission filter 640-700 nm).

### Evaluation of mitochondrial ROS level in live FDB myofibers

The fluorescent dye MitoSOX™ Red (M36008, Invitrogen) was used to evaluate mitochondrial superoxide level in live FDB myofibers. FDB myofibers were incubated with 1 μM MitoSOX™ Red in Krebs solution for 10 min at 37°C. The fluorescence intensity of MitoSOX™ Red was recorded on Leica TCS SP8 confocal microscope (Leica Microsystems Inc., Germany). MitoSOX™ Red was excited by 514 nm laser (emission filter 570-600 nm). All parameters for imaging collection were kept the same between control and experimental groups, which includes the power of the laser used, the pinhole, and the gain of the fluorescence recording.

### Evaluation of membrane repair function in live FDB myofibers

We adapted a laser-induced membrane damage protocol from our early publications[23, 29] to evaluate the membrane repair function of live myofibers. 2 μM of FM 1-43 fluorescent dye were added to the medium of freshly isolated FDB myofibers[23, 26] with 50 μM n-benzyl-p-toluene sulfonamide to prevent myofiber contraction[20, 35]. Using the FRAP protocol on the Leica TCS SP8 confocal microscope, a small area of the myofiber (12 μm X 12 μm) was exposed to a high intensity laser (488 nm, 50% power) to cause a localized muscle membrane injury. The time-dependent accumulation of FM 1-43 fluorescent signal inside the myofiber after the laser-induced membrane injury was recorded every 10 s for 5 min. FM 1-43 was excited by 488 nm laser (emission filter 550-610 nm). All parameters for imaging collection were kept the same for all tested myofibers, which includes the power of the laser used, the pinhole, and the gain of the fluorescence detector. The individual data point of the fluorescent intensity was calculated as (F-F_0_), in which the background fluorescence (F_0_) was corrected for each data point.

### Quantification of membrane integrity of diaphragm muscle

EB dye (1% in PBS, 10μl/g body weight) was applied to mice via IP injection[29]. 16 hours later, the live diaphragm muscle was removed from the mice and immediately immersed in Krebs solution for evaluation of the EB fluorescence intensity retained inside myofibers under a confocal microscope. EB was excited by 633 nm laser (emission filter 640-700 nm). All parameters for imaging collection were kept the same between control and experimental groups, which includes the power of the laser used, the pinhole, and the gain of the fluorescence recording.

### Immunohistochemistry of mouse and human muscle

For whole-mount fixed muscle assay, the intact diaphragm, TA, EDL, and soleus muscles were dissected from the experimental mice and fixed in either methanol (precooled at −20 °C) for 10 min or 4% paraformaldehyde (PFA) overnight at 4 °C. For PFA fixed samples, the reaction was stopped by PBS containing 1% glycine. The samples were then washed with PBS, dehydrated and rehydrated through a graded series of alcohol. Pre-blocking was done at room temperature for 2 hours in blocking buffer containing 3% goat serum, 3% BSA, 0.1% Tween-20, 0.1% Triton X100, and 0.1% NaN_3_. The whole-mount fixed muscle samples were then incubated with primary antibody at 4 °C overnight (anti-neurofilament: Abcam, ab8135 1:250 dilution; anti-Mg53 antibody: 1:200 dilution [32, 36–38]). After PBS washing, the samples were incubated for 4 hours with Alexa Fluor conjugated secondary antibodies of the corresponding species/Isotype (1:1000). The samples were then washed again, cleared in glycerol and mounted in anti-fade mounting media (Tris buffer containing 60% glycerol and 0.5% N- propyl gallate) for imaging under confocal microscope. For the NMJ staining, BTX-Alexa Fluor 488 (1:1000 dilution) was added during the 2^nd^antibodies’ incubation.

Formalin-fixed, paraffin-embedded slides of post-mortem ALS and non-neurological control decedent muscle were obtained from the *Target ALS Multicenter Postmortem Tissue Core*. All decedents underwent standard autopsies with consents for autopsy obtained by next-of-kin after death, and HIPAA Form 5 exemptions to access and share de-identified clinical data from decedents. Slides were provided blinded to whether they were from ALS or control autopsies, and then unblinded after staining and analysis was completed.

For paraffin sectioning, muscle samples were fixed in 4% paraformaldehyde (PFA) overnight at 4 °C, washed with PBS, and dehydrated through a graded series of alcohol. Clearance was done with xylene and paraffin embedding was carried out under negative pressure for 3 hours. Samples were cut into 8μm sections for immunostaining. Antigen retrieval was carried out at 95 °C for 20 min in citrate buffer (pH 6.0).

For dissociated FDB myofibers, the samples were allowed to attach to laminin (sc-29012) coated glass bottom dishes for 30 minutes in culture media containing Alexa Fluor coupled BTX (1:1000) before fixation (4% PFA for 15 minutes at 37 °C). The pre-blocking, primary, and secondary antibody incubation steps were the same as described above.

### Immunoblotting assay

The mouse serum samples (1.5 μl each) were resolved in 8.7% SDS polyacrylamide gels and transferred to a polyvinylidene difluoride (PVDF) membrane and probed with an anti-MG53 antibody 1:2000. Ponceau S staining was used to verify equal loading of the serum samples. Proteins from mouse TA muscle were extracted with RIPA/protease inhibitors and resolved by 10% SDS-PAGE, then transferred to PVDF membrane and probed with antibodies against MG53 antibody 1:5000, and GAPDH (1:10,000, from CST). Protein bands were visualized with ECL reagents under ChemiDoc Imaging system (Bio-Rad Laboratory). Band intensity was analyzed with ImageJ software (NIH, Bethesda, MD).

### Evaluation of serum creatine kinase (CK) activity

Blood samples (~100μl/mouse) were collected from the tail vein of mice at rest. Then the same group of mice were subjected to a 30 min running protocol at 18 m/min, 15° downhill. Right after completion of the running protocol, the blood samples were collected again. The serum was collected from supernatant after centrifuging the blood samples at 2000g for 10 min at 4° C. Freshly made serum samples were used to measure CK activity following the protocol provided by Sigma (MAK-116). Briefly, for each reaction, 10 μl of serum was mixed with 100ul assay buffer, the mixture was then loaded onto a 96 well plate for recording the absorbance at 340nm at time intervals of 5 min up to 40 min in a plate reader (SpectraMax i3x). The CK activity was calculated according to the formula provided:

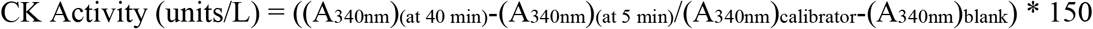

### Quantification of motor neurons in lumbar spinal cord with Nissl Staining

The lumbar portion of the spinal cord was collected from the mice and fixed in 3.7% PFA at 4°C overnight and stored in 70% ethanol until use. After embedding in 6% LMP agarose, the fixed spinal cords were cut in 25μm sections starting at the upper part to subsequently collect 20 sections from each lumbar spinal cord using a Vibratome (Leica VT1000S, Germany). The spinal cord sections were stained with 0.1% Cresyl violet acetate. The motor neurons (Nissl substrate-positive with diameter lager than 25 μm) in the anterior horn region of each section were quantified in a double-blind manner for different treatment groups.

### Confocal imaging and image analysis

Leica TCS SP8 confocal microscope (Leica Microsystems Inc., Germany) was used for imaging. Images were captured with either 40X, 1.2 NA water immersion objective or 63X, 1.4 NA oil immersion objective. Imaging was conducted at room temperature (~23°C). Background correction, binary segmentation, polyline kymograph analysis (for fluorescent intensity profiling over distance) were performed using Fiji (ImageJ). For mouse movement tracking, the video files were registered using ImageJ (Template Matching Plugin) and tracking analysis was conducted using “Manual Tracking with TrackMate” plugin. Individual mouse was tracked by creating spots (with diameter equivalent to mouse belly width) at the center of the torso in each frame. Percentage of time showing constrained movement was defined as the percentage of the recording time in which the mouse moved at less than 1 cm/sec speed for 3 seconds or longer.

### PEGylation of rhMG53

PEGylation is a well-established method for increasing the circulating half-life of therapeutic proteins[39]. We conducted a study with mPEG-SVA (purchased from LaySan Bio, Inc.) modification of rhMG53. 40mg rhMG53 protein was dissolved in PBS (pH = 8) solution at 1 mg/ml concentration in 4 °C. 40 mg PEG-SVA was added into the rhMG53 solution and mixed gently. The reaction of PEGylation was carried out at 4 °C overnight. Un-conjugated PEG-SVA was filtered out by Amicon 30 ultrafilter with PBS. PEG-rhMG53 was stored in −20 °C for long-term storage or 4 °C for short-term usage and to avoid the freeze thaw cycle.

### Pharmacokinetic evaluation of PEG-rhMG53 in rats

Rats were anesthetized by isoflurane inhalation. rhMG53 or PEG-rhMG53 were injected intravenously via tail-vein. Blood samples were collected from the tail-vein at 15 min, 30 min, 1 hr, 2 hr, 6 hr, 12 hr, 24 hr, 48 hr (rhMG53 only), and 96 hr (PEG-rhMG53 only) respectively. Serum was collected by centrifugation of clogged blood at 8000 rpm at 4 °C for 10 min and diluted at 1:50 for ELISA assay to quantify the serum levels of rhMG53. Specifically, the ELISA plate (Nunc-Immuno™ plates, 96 well-plate, MaxiSorp) was coated with 100ul/well an anti-MG53 rabbit monoclonal antibody (10 ug/ml) in coating buffer (Na_2_CO_3_: 3.18 g; NaHCO_3_ 5.88g to 1000ml, pH=9.6), at 4 °C, overnight. ELISA washing and blocking reagent were purchased from KPL. Diluted serum samples as well as rhMG53 standards were added into the coated ELISA plates and incubated for 1.5 hrs. After 4X washing, biotinylated anti-MG53 5259 (1:500, 100ul/well) was added as the detection antibody and incubated for 1.5 hrs. After 4X washing, HRP-Streptavidin (HRP-Conjugated Streptavidin (Thermo Scientific, Cat No: N100), 1:5000 dilute in blocking buffer) was added and incubated for 30 min at room temperature. After 5X washing, KPL SureBlue Reserve™ TMB microwell peroxidase substrate (Cat No: 53-00-02) was added and incubated till blue color develops, then read O.D. at 650 nm. Note that the reason we used rats instead of mice for the pharmacokinetic evaluation of PEG-rhMG53 is because rat has much more total blood volume (25 ml) than mice (1.5 ml). Thus, the multiple blood collections would have minimal effect on rats.

### Intravenous administration of rhMG53 and PEG-rhMG53

Stock solution of rhMG53 or PEG-rhMG53 was prepared by dissolving rhMG53 or PEG-rhMG53 in sterilized saline solution at the concentration of 2 mg/ml. The 3/10 cc insulin syringe with gauge 31 needle was used for the injection. The experimental G93A mouse received isoflurane inhalation (1-2%) to reach an appropriate anesthesia state, at which time the mouse showed no response to pinching at their toes. Then, submandibular vein area of the mouse was cleaned with 70% alcohol wipe, and then rhMG53 or PEG-rhMG53 solution (~30 μl) was injected into the submandibular vein. The G93A mice in the control group received same amount of saline injection. The isoflurane inhalation was immediately stopped after the injection. Usually, the mouse resumed normal activities within five minutes after the stop of isoflurane inhalation. Submandibular vein on both sides was used alternatively for the intravenous administration.

### Numeric data presentation and statistics

All measurements were taken from distinct samples. Data were presented as mean ± S.E. of the independent determinations except **Fig. 5g**, which used median ± MAD (median absolute deviation) due to the relatively large variation of the dataset, and the survival test in **Supplementary Table 3**. Statistical comparisons were done using students’ t-test (two-sided) for single mean or ANOVA test for multiple means when appropriate (we assume data to be normally distributed). Pearson’s Chi-squared tests were carried out by RStudio. Pearson’s correlation coefficient for colocalization analysis was calculated by Coloc 2 plugin in ImageJ following the equation below:

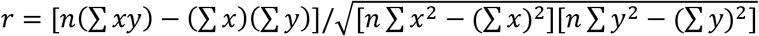

Line plots were generated in either Sigmaplot (Systat Software Inc.) or Excel. Box-and-dot plots were created by the ggplot2 package of RStudio. The box bottom, median line, and box top represent the 25th (Q1), 50th (Q2) and 75th percentile (Q3), respectively. Whisker ends represent Q1 – 1.5*IQR and Q3 + 1.5*IQR, respectively. IQR is interquartile range (Q3-Q1). P < 0.05 was considered statistically significant.

## Results

### Increased susceptibility to diaphragm membrane injury is an early finding in SOD1(G93A) mice

Transgenic mice expressing the human SOD1^*G93A*^ mutation (G93A) are a highly characterized and widely used model for preclinical investigation of pathogenic mechanisms in ALS[40, 41]. As illustrated in **Fig. 1a**, we performed intraperitoneal (IP) injections of Evans blue (EB) dye to wild type (WT) and G93A littermates at 2 months of age (pre-symptomatic stage, prior to ALS onset)[34]. We harvested the diaphragm muscles 16 hours later, immediately after the mice performed a 30 min running protocol (18 m/min, 15° downhill), and evaluated EB retention within muscle myofibers. The amount of EB fluorescence measures the extent of muscle membrane leakage. The G93A diaphragm muscles displayed significantly higher EB levels under resting condition compared to WT (**Fig. 1b**). The 30 min downhill protocol further dramatically enhanced EB accumulation in the G93A diaphragm (**Fig. 1c**), but there was no change in the WT mice. The same exercise-induced excessive membrane damage was also observed in the tibialis anterior (TA) muscle derived from the G93A mice (**Supplementary Fig. 1**). Measurement of serum creatine kinase (CK) showed significant elevation in G93A mice after down-hill running (**Supplementary Fig. 1d**).

**Fig. 1.**
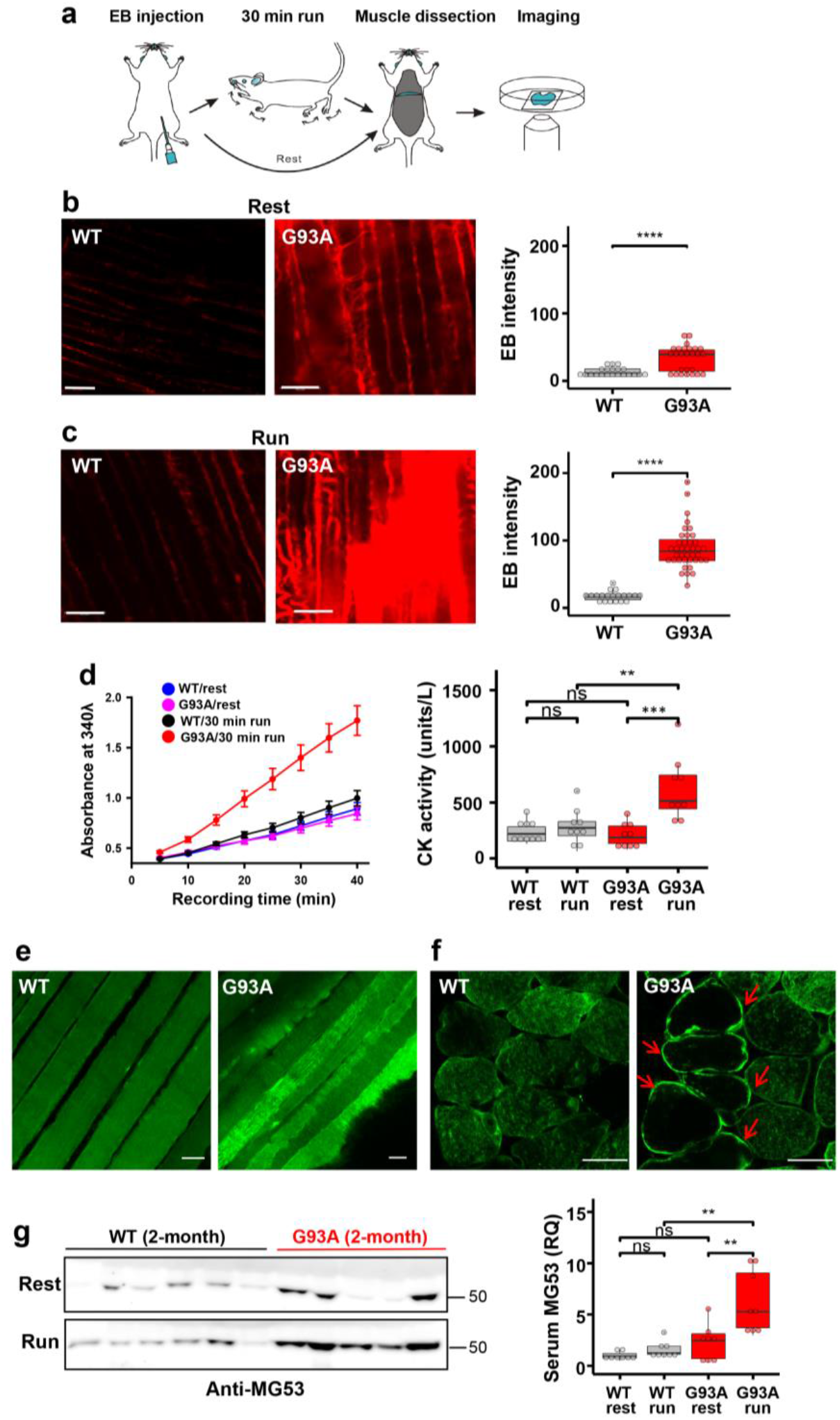
Diaphragm susceptibility to membrane injury is an early pathological event in ALS. (**a**) Schematic diagram for evaluation of diaphragm integrity in ALS mice. (**b**) Compared with WT mice, diaphragm muscle derived from G93A littermates (2-month old) showed increased EB intensity without running (WT: 12.8 ± 1.5, n = 21 from 4 mice; G93A: 33.4 ± 3.7, n = 26 from 2 mice, **** P < 0.0001). Scale bars: 20 μm. (**c**) 30 min downhill running caused drastic elevation of EB in diaphragm derived from the G93A mice (WT17.30 ± 1.69, n = 20 from 3 mice; G93A: 88.49 ± 4.92, n = 39 from 4 mice, **** P < 0.0001). Scale bars: 20 μm. (**d**) Quantification of CK activity in serum samples derived from G93A and WT littermate mice (2-month old) before and after 30 min running. n = 9 for each group, **P < 0.01, ***P < 0.001, ns: not significant. ***Left panel*** shows the time-dependent reading of the absorbance (at 340 nm) of serum samples in a 96-well plate; ***Right panel*** shows the calculated CK activity derived from the absorbance reading (see Method Section). (**e**) Immunostaining revealed homogenous patterns of MG53 in WT diaphragm, whereas the G93A diaphragm showed aggregation and patching patterns of MG53 after running, indicative of membrane injury. Scale bars: 20 μm. (**f**) Cross sectional staining revealed “ring-like” patterns of MG53 in G93A diaphragm (*arrows*), which were less frequent in WT diaphragm. Scale bars: 20 μm. (**g**) Immunoblotting of serum MG53 levels in WT and G93A littermate mice (2-month old) before (rest) and after 30 min running (run). Ponceau S staining indicated equal loading of the serum samples (see **Supplementary Fig. 2**). WT mice showed no significant changes in the serum MG53 level before and after running (WT/rest: 1.00 ± 0.14 *vs* WT/run: 1.58 ± 0.28, ns: P > 0.05). After running, G93A mice showed significant increase of the serum MG53 level (G93A/rest 2.29 ± 0.63 *vs* G93A/run: 6.21 ± 1.09, **P < 0.01). n = 8/group.

We performed MG53 immunostaining in diaphragms derived from WT and G93A mice immediately after running. The whole-mount preparation (longitudinal view) (**Fig. 1e**) revealed distinct segments with increased MG53 in the G93A diaphragm, which were rarely seen in the WT diaphragm. In transverse sections (**Fig. 1f**), the increased MG53 staining clearly localized to the sarcolemma in some G93A myofibers (*red arrows*), in contrast to the predominantly intracellular distribution of MG53 in WT myofibers. This observation is consistent with the notion that G93A diaphragm muscles are susceptible to exercise-induced injury, and endogenous MG53 localizes to the areas of membrane damage in ALS muscle, consistent with its known functions in promoting plasma membrane repair.

Immunoblotting showed elevated MG53 levels in sera derived from the G93A mice after running (**Fig. 1g** and **Supplementary Fig. 2**), which correlates with the increases in serum CK levels (**Fig.1d**). A similar correlation between serum CK and MG53 measurements was reported in *mdx* mice, a model of Duchene muscular dystrophy with compromised muscle membrane repair[29]. Since these striking diaphragm and tibialis anterior myofiber injuries in the G93A mice (2-month old) occur prior to the onset of ALS symptoms, these findings suggest that fragility of skeletal muscle represents an early pathological event in ALS.

### NMJ is an active site of injury-repair by MG53 under physiological conditions that is lost in ALS

We showed previously that segmented mitochondrial defects appeared locally at NMJs in the G93A mice[20, 21]. This prompted us to examine whether the NMJ itself is an active site of contraction-induced membrane injury, and if MG53 plays a role in repair of NMJ injury. Immediately after the 30 min running, freshly isolated flexor digitorum brevis (FDB) myofibers from 2-month old mice were fixed for staining with α-Bungarotoxin (BTX) to mark NMJ and anti-MG53 antibodies. We found that MG53 accumulated at the NMJ area and formed patches in WT myofibers (n = 130, 4 mice) (**Fig. 2a**, *left*), suggesting that MG53 contributes to the maintenance of NMJ integrity under physiologic conditions. While MG53 also accumulated at the NMJ area in G93A myofibers (n = 109, 3 mice), there were also separate intracellular MG53 aggregates in proximity to NMJs in about 10% of G93A myofibers (**Fig. 2a**, *right,* see also **Supplementary Fig. 3**). Moreover, FDB myofibers derived from the G93A mice at an advanced stage of ALS (4-month old) showed increased abnormal MG53 aggregation near NMJs even without running (**Fig. 2b** and **Supplementary Fig. 4**). Such MG53 aggregation suggests potential impaired tissue-repair capacity that may manifest into NMJ degeneration during ALS disease progression.

Using the entry of a cell-impermeable fluorescent dye FM 1-43 as a measure of muscle myofiber integrity[23, 42], we tested the possibility that ALS-associated muscle injury originates focally at NMJs. **Fig. 2c** shows an overlay of FM 1-43 and BTX, indicating FM 1-43 enriches at NMJ when applied to the medium. In WT myofibers, FM 1-43 was restricted to the NMJ and there were barely any FM 1-43 signals inside myofibers after a 20 min dye incubation (**Fig. 2d**). In contrast, 59% ± 7% of FDB myofibers from three 4-month old G93A mice showed intracellular uptake of FM 1-43 dye, which formed a gradient centered around the NMJ (**Fig. 2e**), indicating that FM 1-43 dye entered the cells preferentially through injured NMJs.

**Fig. 2.**
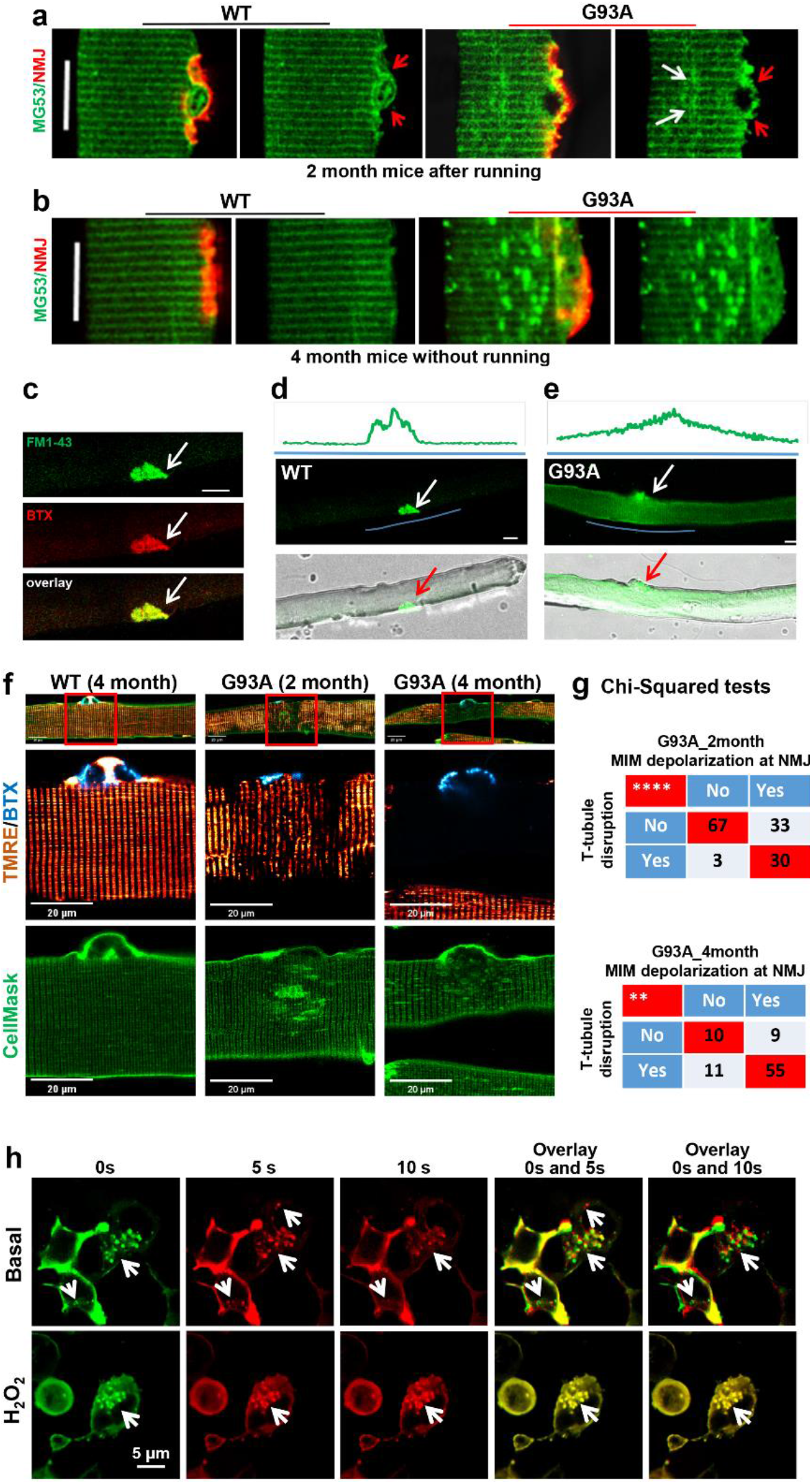
Disruption of physiological injury-repair process by MG53 at NMJ upon ALS progression. (**a**) FDB myofibers, obtained from WT and G93A mice (2-month old) subjected to 30 min running, were co-stained with MG53 antibody (green) and BTX (red). MG53 formed patches covering the NMJ site in both WT and G93A myofibers. However, intracellular MG53 aggregates start to appear near the site of NMJ in the G93A myofibers. Scale bars: 20 μm. (**b**) FDB myofiber derived from G93A mice at the advanced stage of ALS (4-month old) displayed extensive MG53 aggregates near NMJ without running. FDB myofiber derived from WT littermates showed a uniform pattern of MG53. Scale bars: 20 μm. (**c**) FM 1-43 and BTX co-localized at the NMJ of FDB myofibers (white arrows). (**d**) No intracellular FM 1-43 fluorescence was observed in the WT myofiber at 20 min after incubation with FM 1-43. (**e**) FM1-43 entered the G93A myofiber via the NMJ region. Green line highlights the region for fluorescence intensity profiling. Scale bars: 20 μm. (**f**) FDB myofibers were loaded with BTX (cyan), TMRE (red) and CellMask (green) simultaneously. At the site of NMJ, mitochondria depolarization (loss of TMRE fluorescence) occurred in myofibers derived from both 2-month and 4-month old G93A mice, accompanied by disrupted T-tubule network (marked by CellMask). Normal mitochondria and organized T-tubule network were detected at the site of NMJ of WT myofiber. Scale bar: 20 μm. (**g**) Chi-Square tests indicate positive correlations between mitochondrial inner membrane (MIM) depolarization and T-tubule disorganization at the site of NMJ of G93A mice at both 2 months and 4 months of age (**P < 0.01, ****P < 0.0001, 3 mice/group). (**h**) The representative time-lapse images of C2C12 cells with overexpression of GFP-MG53 in the presence and absence (basal) of 1 mM H_2_O_2_. The overlay images were generated using images at 0s and 5s, or 0s and 10s. White arrows indicate GFP-MG53 vesicles.

We next examined whether exercise could exacerbate membrane injury at NMJs of G93A muscle. For this purpose, three pairs of 2-month old WT and G93A littermate mice performed the running protocol for 30 minutes. 72% ± 2% of G93A FDB myofibers showed NMJ-centered intracellular FM 1-43 gradient, whereas only 6% ± 2% WT myofibers exhibited this phenomenon (P < 0.001). These data substantiate the finding that NMJs are focally more susceptible to injury than other regions of the sarcolemma, and exercise-induced NMJ injury is dramatically exacerbated in the G93A ALS mice.

In skeletal muscle, the cell surface membrane travels deep into the myofibril forming transverse tubules (T-tubules). Freshly isolated FDB myofibers were incubated with BTX, CellMask, and TMRE to locate the NMJs, T-tubule network and record mitochondrial potentials, respectively (**Fig. 2f**). Normal polarized mitochondria and organized T-tubule networks were observed in WT myofibers (**Fig. 2f**, *left*). Myofibers derived from the 4-month old G93A mice showed fully depolarized mitochondria at NMJs (**Fig. 2f**, *right*), and those derived from the 2-month old G93A mice showed partially depolarized mitochondria (**Fig. 2f**, *middle*). Remarkably, disorganized T-tubule networks were observed near the NMJs in both 2- and 4-month old G93A myofibers, including regional loss of T-tubules (hollow area under the NMJ) and formation of enlarged spheres. Pearson’s Chi-squared tests demonstrated a significant correlation between T-tubule disruption and mitochondrial depolarization at the NMJ in both 2-month (*P* < 0.0001) and 4-month (*P* < 0.01) G93A myofibers (**Fig. 2g**). These data confirm that mitochondrial dysfunction is associated with the disruption of cell membrane integrity at NMJs of the ALS muscle.

We and others have shown that mitochondrial dysfunction is associated with enhanced reactive oxygen species (ROS) production in the ALS mouse muscle[14, 18, 43], which could impact the intrinsic membrane repair function of MG53[44, 45]. Using our established live cell imaging method, we examined whether oxidative stress impact the traffic of MG53 vesicles inside C2C12 cells overexpressing GFP-MG53[46]. The 2D x-y time-lapse images were continuously recorded for 10 seconds in the presence or absence (basal) of 1 mM H_2_O_2_. Representative images at 0 sec (0s, pseudo color green), 5 sec (5s, pseudo color red) and 10 sec (10s, pseudo color red) were selected for generating the overlay images (**Fig. 2h**). The overlay images of 0s with 5s or 0s with 10s provide a visualization of GFP-MG53 vesicle dynamics. In the overlay images, non-moving vesicles are marked by yellow color (completely overlap), while moving vesicles are indicated by the red and green colors (no overlap). Note that there is almost no detectable movement of GFP-MG53 vesicles in the presence of H_2_O_2_, while moving GFP-MG53 vesicles were detected under basal condition.

### Impaired MG53 membrane repair function is a common pathological feature of ALS muscle

With the support of the *Target ALS Human Postmortem Tissue Core*, we obtained paraffin-embedded diaphragm and psoas muscle sections from both sporadic and familial ALS decedents, and non-ALS controls (**Supplementary Table 1** and **Supplementary Table 2**). Both longitudinal and transverse sections of human ALS diaphragm and psoas muscle showed dramatic abnormal sarcolemmal and intracellular MG53 aggregates. In contrast, the muscle samples of non-neurological control decedents showed only a few scattered MG53 aggregates (**Fig. 3a** and **Supplementary Fig. 5**). It is not unexpected to see a few MG53 aggregates in non-ALS muscle, as MG53-mediated membrane repair also occurs in normal conditions, but to a much lesser extent. As shown in **Fig. 3b**, this same staining pattern was observed in longitudinal (**Fig. 3b**, *left*) and transverse sections (**Fig. 3b**, *right*) of diaphragm muscle from the 4-month old G93A mice. The abnormal intracellular MG53 aggregation was observed in all 4-month old G93A muscles examined, including extensor digitorum longus (EDL), soleus, and tibialis anterior (TA) (**Supplementary Fig. 6a**). The data from both familial and sporadic ALS decedents and the G93A mouse model suggest that compromised MG53-mediated muscle membrane repair function could be a common pathology in ALS.

**Fig. 3.**
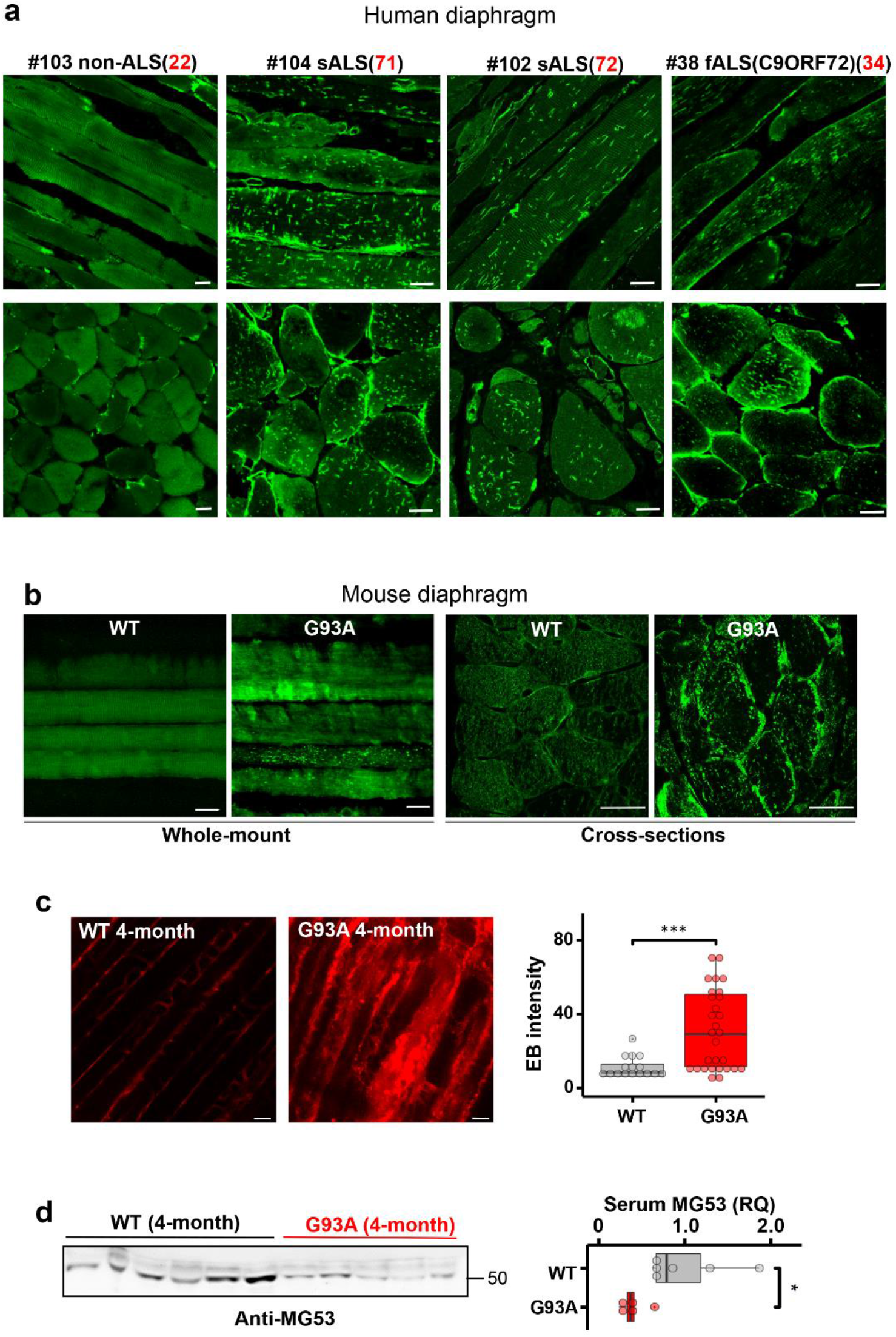
Impaired MG53 membrane repair function is a common pathological feature of ALS. (**a**) Longitudinal (top panels) and cross-sectional (bottom panels) staining of MG53 demonstrated uniform patterns in non-ALS human diaphragm (left); whereas human ALS diaphragms (sALS: sporadic ALS: fALS: familial ALS) displayed extensive intracellular aggregates of MG53. The age at the death was indicated in red fonts. Scale bars: 20 μm. (**b**) Diaphragm muscle derived from a 4-month old G93A mouse (post-ALS onset) also showed intracellular aggregates of MG53. (**c**) Evaluation of the diaphragm muscle cell membrane integrity using EB dye in 4-month old G93A and WT mice at rest. The G93A diaphragm showed enhanced EB accumulation (G93A: 31.1 ± 4.0, n = 29 from 3 mice vs WT: 10.9 ± 1.6, n = 16 from 2 mice, ***P < 0.001). Scale bars: 20 μm. (**d**) G93A mice at the age of 4 months without running showed significantly lower levels of the serum MG53 compared with WT littermates (G93A, n = 5; WT, n = 6, *P < 0.05). Scale bars: 20 μm.

Enhanced EB retention in the diaphragm (**Fig. 3c**) and TA muscles (**Supplementary Fig. 6b**) was also observed in 4-month old G93A mice without running, indicating that muscle injury and fragility continue during ALS progression. Interestingly, the serum level of MG53 in the G93A mice at this late stage of ALS was lower compared with the WT littermates under resting conditions (**Fig. 3d**). As MG53 is a muscle-derived protein, the reduced serum MG53 level likely reflect the progressive loss of muscle mass. It is also possible that the formation of intracellular MG53 aggregates sequesters MG53, limiting its release into circulation. Nevertheless, these findings suggest that endogenous MG53 no longer can sustain the membrane repair function as ALS progresses.

### Recombinant human (rhMG53) preserves the membrane integrity of diaphragm in G93A mice

Using an established protocol for evaluating the muscle cell membrane repair function following laser-induced cell membrane damage[23, 42], we conducted *in vitro* studies to test (1) whether G93A myofibers showed higher laser-induced fragility compared to WT myofibers, and (2) whether exogenous administration of rhMG53 could improve membrane integrity of G93A myofibers. A small area of the FDB myofiber (12μm X 12μm) was exposed to a high intensity UV laser to cause localized cell membrane injury allowing the entry of FM 1-43 dye. As shown in **Fig. 4a**, G93A myofibers (4-month old) showed more intracellular accumulation of FM 1-43 compared with WT, indicating fragility and impaired cell membrane repair mechanism. 10 μg/ml of rhMG53 applied to the culture medium reduced the intracellular accumulation of FM 1-43 in the G93A myofibers. Application of the same amount of bovine serum albumin (BSA) was ineffective. The time-dependent FM 1-43 accumulation following laser-induced injury is shown in **Fig. 4b**, demonstrating that extracellular application of rhMG53 improved the membrane repair function of the G93A muscle. Furthermore, G93A myofibers showed reduced mitochondrial ROS production after incubation with rhMG53 (2 μg/ml) in the medium for 12 hours (**Fig. 4c**).

**Fig. 4.**
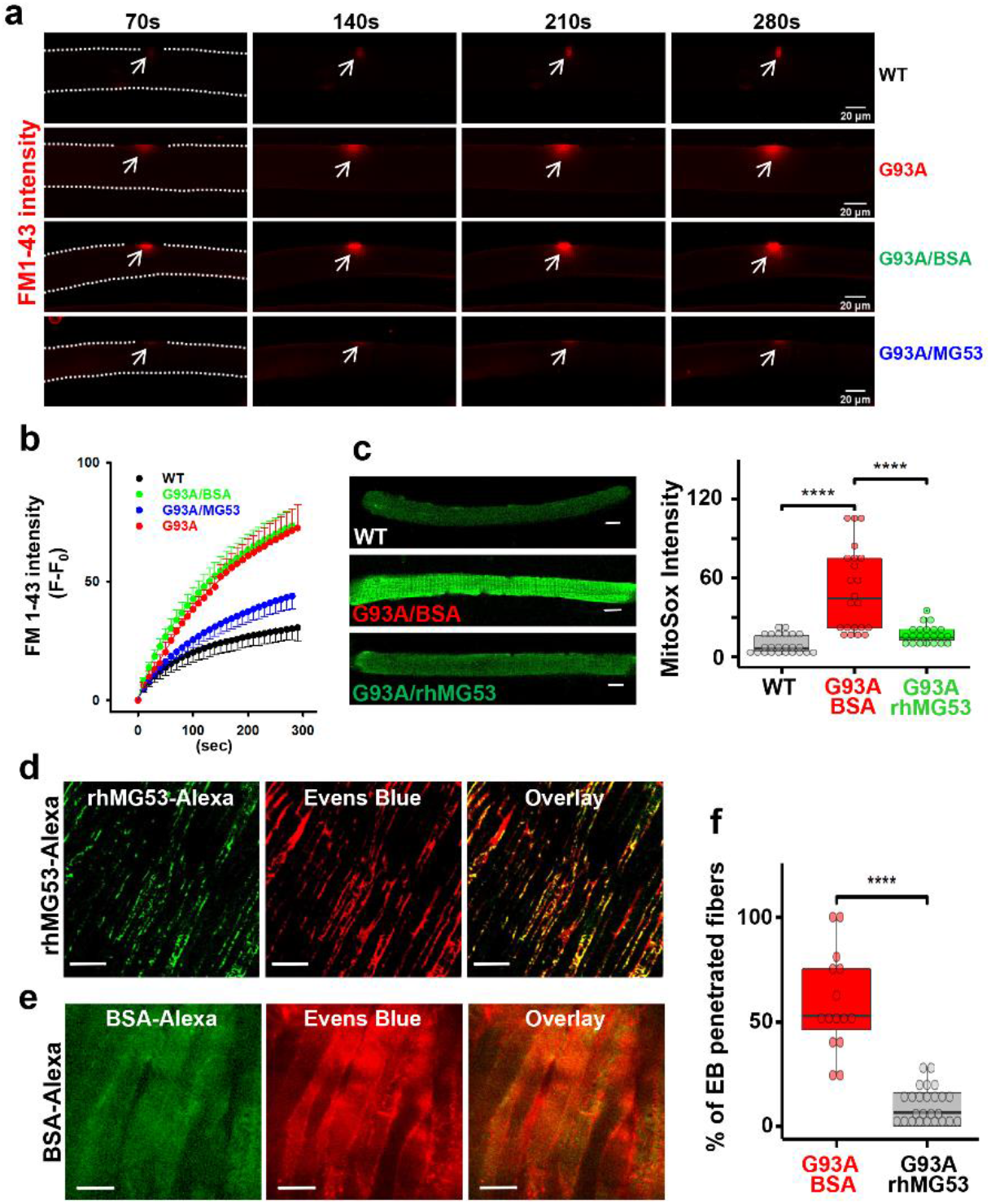
rhMG53 preserved membrane integrity of the diaphragm in G93A mice. (**a**) Time-lapse recordings of FM 1-43 accumulation in FDB myofibers after laser-induced membrane damage. The dashed lines highlight the myofiber outline. (**b**) Plotting of FM 1-43 intensity inside FDB myofibers over time after laser-induced membrane injury. Between G93A and WT (n = 6-9 myofibers/group; P < 0.001), and between G93A/BSA (9 myofibers) and G93A/rhMG53 (10 myofibers) (P < 0.001). (**c**) Live WT and G93A FDB myofibers pre-treated with rhMG53 (2 μg/ml) or BSA (2 μg/ml, as control) were loaded with MitoSox Red for evaluating the mitochondrial superoxide production. rhMG53 treatment significantly reduced mitochondrial superoxide level in G93A FDB myofibers (WT, n = 24; G93A+BSA, n = 22; G93A + rhMG53, n = 22, ****P < 0.0001). Scale bars: 20 μm. (**d**) Exercise-induced accumulation of EB in diaphragm derived from G93A mice (2-month old) was reduced with IV administration of rhMG53-Alexa (2 mg/kg). rhMG53-Alexa targeted to the sarcolemma of the diaphragm. Scale bars: 20 μm. (**e**) Diaphragm derived from G93A mice receiving BSA-Alexa showed extensive accumulation of EB. BSA-Alexa did not target to the sarcolemma. Scale bars: 20 μm. (**f**) Percentage of myofibers with EB penetration was significantly reduced with administration of rhMG53-Alexa (n = 14 for BSA-Alexa488, n = 25 for rhMG53-Alexa488 from 3 pair of G93A mice, ****P < 0.0001).

We next tested whether intravenous (IV) administration of 2 mg/kg rhMG53-Alexa (or BSA-Alexa as a control) in the G93A mice - immediately prior to running, attenuates diaphragm membrane injury. The dosage of rhMG53 was determined based on our previous publications with rhMG53 in preservation of membrane integrity in multiple organs[30, 32, 33]. After running, diaphragm muscles were immediately collected for live cell imaging of Alexa and EB simultaneously. As shown in **Fig. 4d**, mice treated with rhMG53-Alexa showed prominent membrane patches of MG53-Alexa, indicating that rhMG53-Alexa targets to injured diaphragm muscles, similar to the previous study with the *mdx* mice[29]. In contrast, diaphragm derived from the G93A mice receiving BSA-Alexa showed a diffuse pattern, indicating that BSA-Alexa could not form membrane patches (**Fig. 4e**). Compared with BSA-Alexa treated mice, the proportion of myofibers positive for intracellular accumulation of EB was significantly reduced in the diaphragm muscles from rhMG53-Alexa treated mice (**Fig. 4f**), confirming the efficacy of rhMG53 to preserve diaphragm membrane integrity *in vivo*.

### Systemic application of rhMG53 preserved NMJ integrity and extended the life span of G93A mice

Our ultimate goal is to develop rhMG53 as a treatment for ALS, which would be initiated after diagnosis – and thus after symptom onset – to ALS patients. Therefore, we evaluated the efficacy of rhMG53 administration to G93A mice after ALS onset by starting the treatment at the age of 3 months[47]. We first adapted a method to quantify NMJ innervation in diaphragm muscle[48] of adult mice. **Fig. 5a** shows a representative image of a mouse diaphragm stained with anti-neurofilament (NF) antibodies (red fluorescence) and BTX (green fluorescence). By changing the focal plane and performing a 3D-scan of a zoomed-in area, we could examine NMJ in the entire diaphragm. The 3D-scan projection allowed us to distinguish well innervated NMJs from partially innervated or denervated NMJ (**Fig. 5b**). Then, ten G93A mice (3-month old) were divided into two cohorts, one received IV injection of rhMG53 (2 mg/kg, daily), and the other received saline for 2 weeks. **Fig. 5c** shows representative 3D-projection images of the diaphragm muscle from rhMG53 and saline treated mice. The innervated NMJ area was defined by the overlapping area between BTX and NF signals. Quantitative analyses of NMJ innervation are presented in **Fig. 5d**, demonstrating that rhMG53 treatment could significantly preserve NMJ integrity, maintain well innervated and partially innervated NMJs, as well as reduce the proportion of denervated NMJs compared with saline-treated controls. In a separated experiment, motor neurons in the anterior horn region (Nissl staining-positive with diameter lager than 25 μm) were counted (**Fig. 5e and 5f**). The 2-week rhMG53 treatment significantly preserved the number of motor neurons in spinal cords from the G93A mice (rhMG53 vs saline, P < 0.01) (**Fig. 5g**), suggesting that this treatment also alleviated motor neuron degeneration.

**Fig. 5.**
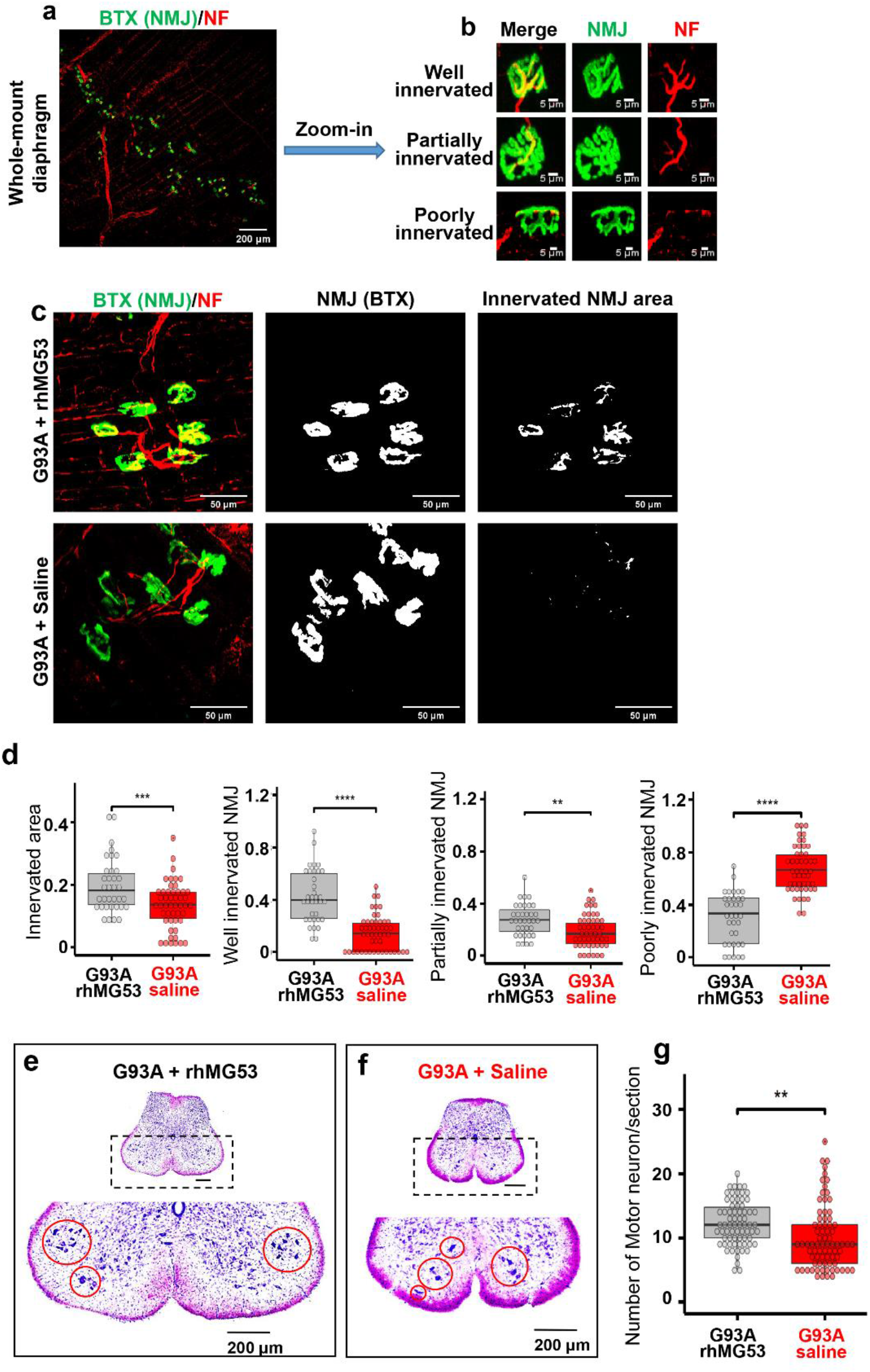
Systemic application of rhMG53 preserved NMJ integrity and promoted motor neuron survival in ALS mice. (**a)** Whole-mount fixed diaphragm muscle stained with the NF antibody (red) for marking axonal terminals and BTX (green) for detecting NMJ. (**b**) Z-stack images of well (the ratio of innervated area >20%), partially (the ratio of innervated area between 10% and 20%), and poorly innervated NMJs (the ratio of innervated area < 10%) in diaphragm muscle. (**c**) Z-stack images of diaphragm muscle from G93A mice receiving 2-week rhMG53 treatment or saline-control (*left panels*). The area of individual NMJ defined by BTX was presented (*central panels*). The innervated area of NMJ was defined by the area overlapping with NF (*right panels*). (**d**) Comparing the ratio of innervated NMJ area in rhMG53 (n = 39, 4 mice) and saline (n = 45, 5 mice) treated diaphragm muscles of G93A littermate mice, as well as the ratio of well, partially, poorly innervated NMJs. rhMG53 treatment significantly preserved the innervation of NMJ in diaphragm muscle of G93A littermate mice. **P < 0.01, ***P < 0.001, ****P < 0.0001. (**e, f**) Images of the lumbar spinal cord section of G93A mice (with 2-weeks of rhMG53 or saline treatment from the age of 3 months). (**g**) The number of surviving motor neurons per section in G93A littermate mice after two-weeks of treatment with rhMG53 (12.2 ± 0.4) or saline (10.1 ± 0.5) (rhMG53, n = 70 spinal cord sections; saline, n = 77 spinal cord sections; 4 pairs of G93A mice per cohort, ** P < 0.01).

Endogenous MG53 is predominantly expressed in striated muscle[23, 49], but not in neurons[33]. Under physiological condition, MG53 in circulation does not cross the brain blood barrier[33]. Therefore, it was unlikely that rhMG53 in circulation directly reached the motor neuron in spinal cord to protect its function. As ALS appears to be a combination of “dying-forward” and/or “dying-back” pathophysiological processes, and the NMJ is the critical site for this bidirectional crosstalk between motor neurons and myofibers, we speculate that the effects of rhMG53 on preserving anterior horn motor neuron cell bodies is likely secondary to the preservation of NMJ integrity, which slows the dying back process. Contributions from a direct effect on motor neurons remain possible.

The short half-life of rhMG53 in circulation (~1 hour) may present a hurdle for treating chronic tissue injuries in ALS[32, 50]. PEGylation is a well-established method for increasing the half-life of therapeutic proteins in circulation[39, 51]. Addition of polyethylene glycol (PEG) often reduces immunogenicity of the PEGylated proteins without a major loss of their biological activity[52–54] and several PEGylated proteins have reached the market[54, 55]. We produced PEGylated rhMG53 (PEG-rhMG53). As shown in **Fig. 6a**, PEGylation of rhMG53 is successful based on the oligomerization pattern of the protein run on SDS-PAGE. We previously developed an *in vitro* assay to evaluate the efficacy of rhMG53 in protecting against membrane damage by measuring LDH in the extracellular solution of culture cells, as LDH leaks from the cell into the extracellular solution following cell membrane injury[25, 26, 29, 56]. The PEG-modification did not affect the membrane repair function of rhM53 as the EC_50_ of LDH release did not change (**Fig. 6b**). Meanwhile the half-life of PEG-rhMG53 increased in circulation from 0.5 to 12 hours (**Fig. 6c**). This elongated half-life in circulation allowed us to test the PEG-rhMG53 in G93A mice by IV injection in every other day for a longer period.

**Fig. 6.**
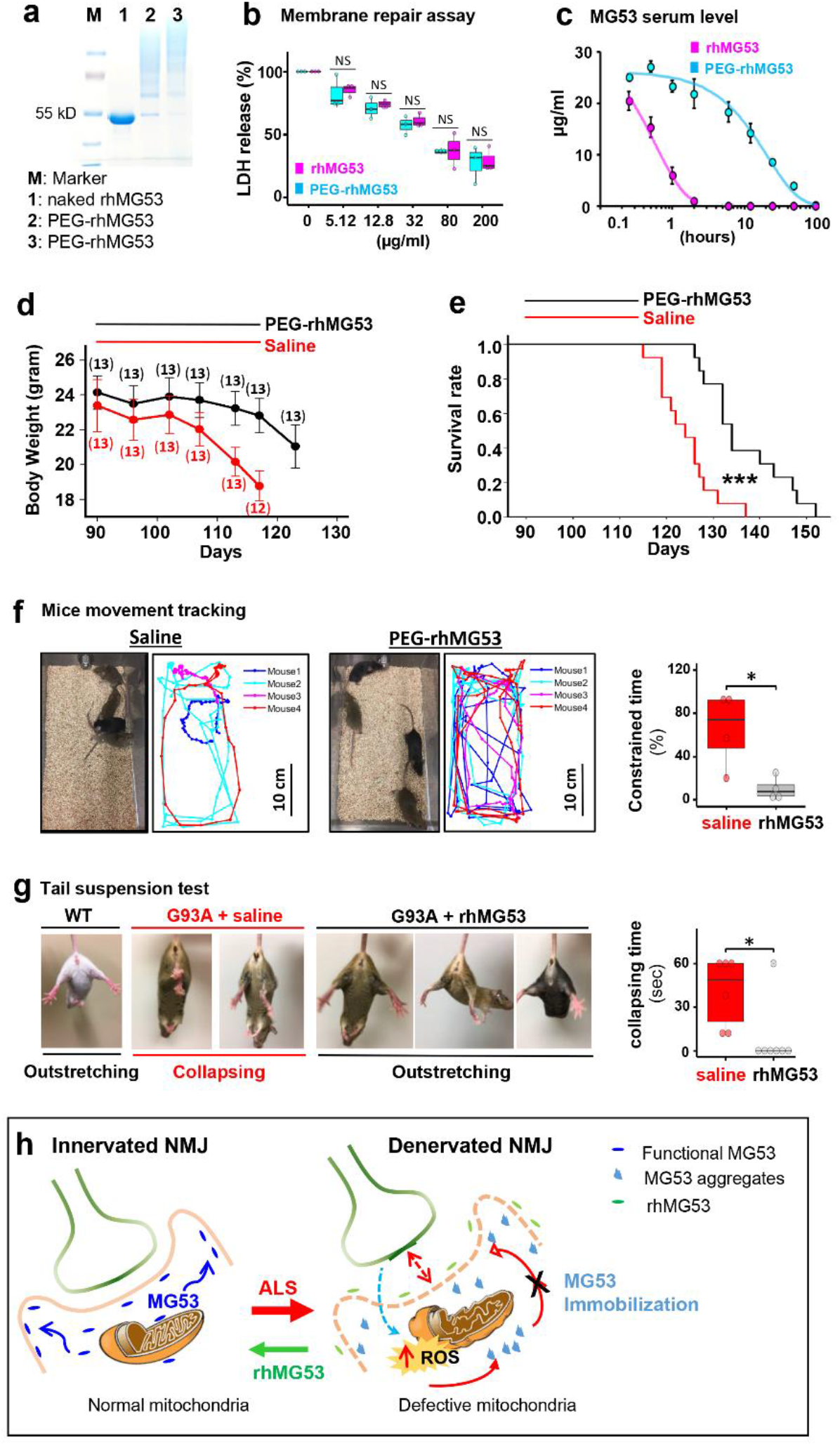
rhMG53 alleviates disease progression and extends the life span of G93A mice. (**a**) PEGylation of rhMG53 is successful based on the oligomerization pattern of the protein running on SDS-PAGE. Lanes 2 and 3 represent two independent rhMG53 PEGylation experiments. (**b**) The chemical modification did not affect the membrane repair function of rhM53 as there were no difference in EC50 of LDH release from the cultured C2C12 cells following a mechanical membrane damage with glass beads between the rhMG53 and PEG-rhMG53 (n = 3 independent experiments). (**c**) Pharmacokinetic (PK) assessment revealed that PEGylation increased serum half-life of rhMG53 in rats from 0.5 hour (rhMG53, n = 4) to 12 hours (PEG-rhMG53, n = 3). (**d**) Bodyweight changes of G93A mice receiving PEG-rhMG53 or saline treatment. The numbers of mice included in each data point were indicated on the plot. (**e**) Survival curve of G93A mice with one-month PEG-rhMG53 or saline treatment. The endpoint (death) of a G93A mouse was defined by the loss of righting reflex within 30 sec when the mouse was place on its side. (n = 13 pairs of G93A littermates, ***P < 0.001). (**f**) Mice movement tracking was performed using 1-minute video recording files of the saline and PEG-rhMG53 treated mice (**Supplementary Movie 1**). Percentage of time showing constrained movement is defined as the percentage of the recording time in which the mouse moved at less than 1 cm/sec speed for 3 sec or longer. (**g**) Representative photos and Collapsing time measurement results of tail suspension tests of G93A mice at the age of 104 day after receiving rhMG53 (n = 7 mice) or saline (n = 6 mice) for 2 weeks. An age-matched WT mouse was included for comparison (* P < 0.05). (**h**) Proposed mechanisms underlying membrane repair defects at NMJ of ALS.

The PEG-rhMG53 (2 mg/kg, IV) was administered to the G93A mice (3-month old) every other day for 30 days. Twenty-six G93A littermate mice (from 3 litters, with both genders included) were divided into two groups, one receiving PEG-rhMG53, and the other receiving saline as a control. The bodyweights of the PEG-rhMG53 or saline treated mice were recorded. One-month of PEG-rhMG53 treatment slowed weight loss compared with the saline treated G93A mice (**Fig. 6d**). Note that the weight loss of PEG-rhMG53-treated mice accelerated again after the treatment ended. This is not unexpected, as the half-life of PEG-rhMG53 is only 12 hours in rodent circulation (**Fig. 6c**). This one-month treatment of PEG-rhMG53 also significantly extended the life span of G93A mice from 124 ± 6 days (saline) to 137 ± 9 days (PEG-rhMG53) (**Fig. 6e**). The endpoint (death) of a G93A mouse was defined by the loss of righting reflex within 30 sec when the mouse was place on its side. The survival days of individual treated G93A mice and the statistics for both genders were listed in **Supplementary Table 3**. While the cohorts were not perfectly gender-balanced per litter, the therapeutic benefits of rhMG53 on survival remained when male and female mice were considered separately (survival increased to 137 ± 7 for male mice and 135 ± 10 for females), when compared to the saline-treated G93A littermate mice. The Chi square test further confirmed that the significant difference between PEG-rhMG53 and saline treated groups is independent of the gender.

**Supplementary Movie 1** illustrates that G93A mice receiving one month of PEG-rhMG53 treatment showed greater mobility compared with saline-treated mice. We tracked the total moving distance of each mouse during one-minute video recordings. The tracking maps indicate that paralysis was more advanced in saline-treated mice, and the time period under constrained motion was significantly shorter in the mice treated with PEG-rhMG53 (**Fig. 6f**). In a separate study, we found that 3-month old G93A mice that received 2-week rhMG53 treatment (2 mg/kg, daily IV injection) showed reduced collapsing time of the hind limbs during the tail suspension test at the age of 104 days (**Fig. 6g**).

## Discussion

We have demonstrated that muscle membrane fragility and damage are increased in ALS G93A mice prior to symptom onset and continue to increase throughout the course of the disease. Furthermore, our study reveals that the site of NMJs are particularly susceptible to injury compared to the rest of the sarcolemma, and MG53 forms membrane patches at NMJ following modest exercise training. Thus, MG53 represents an important physiologic component of protection against injury to the NMJ. Systemic administration of exogenous rhMG53 has beneficial effects to restore diaphragm muscle repair and NMJ integrity and improved the life span of the G93A mice. Abnormal intracellular aggregation of MG53 was observed in multiple types of muscles from the G93A mice. Similarly, the abnormal aggregates were seen in muscle samples from human ALS decedents with both sporadic and familial forms of ALS.

While transient intracellular oxidation initiates MG53-vesicles to form repair-patches[23], exposure of cells to sustained oxidative stress leads to immobilization of MG53’s membrane repair function[44]. It is known that mitochondrial dysfunction in ALS causes intracellular oxidative stress, thus elevated ROS could impede normal MG53 movement to areas of sarcolemmal damage, resulting in the observed aggregation and loss of its tissue repair function. A proposed mechanism underlying the membrane repair defects at the NMJ of ALS is illustrated in **Fig. 6h**. Since NMJ is an active site of neuron-muscle crosstalk, it is conceivable that membrane repair defects initiate from NMJ. MG53 plays a critical role in maintaining the integrity of NMJ and the muscle membrane. During ALS progression, mitochondrial dysfunction causes elevated ROS production, leading to ectopic MG53 aggregation and disruption of MG53’s tissue repair function, which could exacerbate NMJ denervation. Oxidative stress, muscle membrane damage and NMJ denervation could form a vicious cycle promoting muscle wasting and neuronal death in ALS. We demonstrate that exogenously administered rhMG53 forms repair-patches and reduces membrane leakage of the ALS diaphragm muscle. It is possible that reduced cell membrane leakage lowers the energy demand and subsequent mitochondrial ROS production, alleviating this vicious cycle to preserve the integrity of NMJ and muscle membrane, which could also slow down the dying-back process of motor neuron degeneration.

IV administration of rhMG53 or PEG-rhMG53 to the G93A mice after disease onset preserved innervation of the diaphragm and prolonged the lifespan. Studies in mice, rats, and dogs reported no observable toxic effects with long-term administration of rhMG53[29, 32, 36]. While we demonstrate that repetitive IV administration of the PEG-rhMG53 has beneficial effects on G93A ALS mice, further studies are still required to establish the safety profile of the chemically modified rhMG53, and to test the therapeutic efficacy in other ALS animal models.

In rodents and humans, MG53 is present at low levels in blood circulation under physiologic conditions. Given that we observed higher circulating levels in the G93A mice than wild-type littermates, and a correlation with serum CK measurements, it is possible that ALS patients also show different levels of circulating MG53. We intend to examine this further in future studies, including the possibilities that (1) a one-time measurement of MG53 early in disease course could be useful as a “prognostic biomarker” of rate of disease progression, (2) a “predictive biomarker” to identify/cohort select patients who might best respond to therapeutic strategies designed to stabilize/repair damage to myofibers, or (3) a pharmacodynamic biomarker to demonstrate therapeutic effect of treatments to preserve myofiber membrane integrity, including exogenously administered PEG-rhMG53.

The effects of exercise training on ALS progression remains controversial[57, 58], and diaphragm pacing was found to be associated with reduced survival in ALS patients with respiratory insufficiency[59–63]. Our study with the ALS mice demonstrates that even modest exercise training leads to increased damage to the diaphragm, which may exacerbate ALS progression. This appears to be due to the severely compromised membrane repair capacity of the ALS muscle. Based on this finding, one should be cautious in designing exercise-related protocols or diaphragm pacing as alternative interventions to mitigate ALS. However, it might suggest that exercise regimens and/or diaphragm pacing in the presence of exogenously administered rhMG53 could be beneficial.

In summary, we have identified defective muscle membrane repair involving MG53 as a pathological change in ALS mice and postmortem human ALS muscle. Muscle membrane injury occurs early in the course of ALS and contributes to disruption of NMJs. Our study provides proof-of-concept data supporting the beneficial effects of rhMG53 in preserving the integrity of NMJ and muscle cell membrane to alleviate ALS progression. Probing the role of mitochondrial ROS production in modulating MG53-mediated cell membrane repair may have broader implications in understanding the basic pathophysiology of ALS.

## Acknowledgments

We greatly appreciate Kathryn Wilsbach, Kathryn Gallo, Brent Harris, and Galam Khan for providing paraffin-embedded human muscle sections from the Johns Hopkins University and Georgetown University sites of the *Target ALS Postmortem Tissue Core* and the *Johns Hopkins ALS Postmortem Core*; Ms. Erin Haggard for language editing of this manuscript.

## Author contributions

J.Y., J.Z., and J.M. conceived and designed the project; J.Z., J.M., L.O., A.L., J.Y., and X.L. wrote the paper; J.Y., A.L., X.L., K.H.P., X.Z., F.Y., Y.X., D.Y., T.T., and J.Z. acquired the data; J.Y., A.L., X.L., L.O., J.M. and J.Z. analyzed and interpreted the data.

## Supplementary Materials

**Supplementary Fig. 1**. Representative images of TA muscle derived from G93A and WT mice (2-month old) with 30 min running that received EB injection 16 hours early.

**Supplementary Fig. 2**. Additional Western blot data for Figure 1G.

**Supplementary Fig. 3**. Additional representative images for Figure 2A.

**Supplementary Fig. 4**. Additional representative images for Figure 2B.

**Supplementary Fig. 5**. Additional images of anti-MG53 immunostaining of ALS and non-ALS human muscle samples.

**Supplementary Fig. 6**. Additional anti-MG53 immunostaining and EB retention results of muscle samples from G93A and WT mice.

**Supplementary Table 1**. Demographics of human decedents.

**Supplementary Table 2**. Biospecimen information for human samples.

**Supplementary Table 3**. The survival days of individual tested G93A mice separated by gender.

**Supplementary Movie 1**. Video of PEG-rhMG53 treated and control G93A mice.

