## supplementary figures and tables for "MG53 slows neuromuscular junction loss and prolongs survival in ALS"

### Supplementary Fig. 1

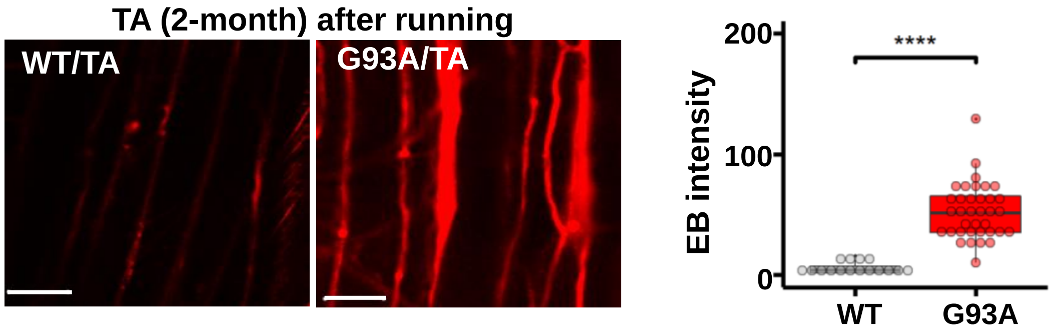

**Supplementary Fig. 1**. Representative images of TA muscle derived from G93A and WT mice (2-month old) with 30 min running that received EB injection 16 hours early. The EB intensity in G93A muscle is significantly enhanced in TA muscle (WT: 5.37 ± 1.13, n = 16 from 2 mice *vs* G93A: 52.40 ± 3.83, n = 36 from 3 mice, ****P<0.0001). Scale bars: 20 µm.

#### Supplementary Fig. 2

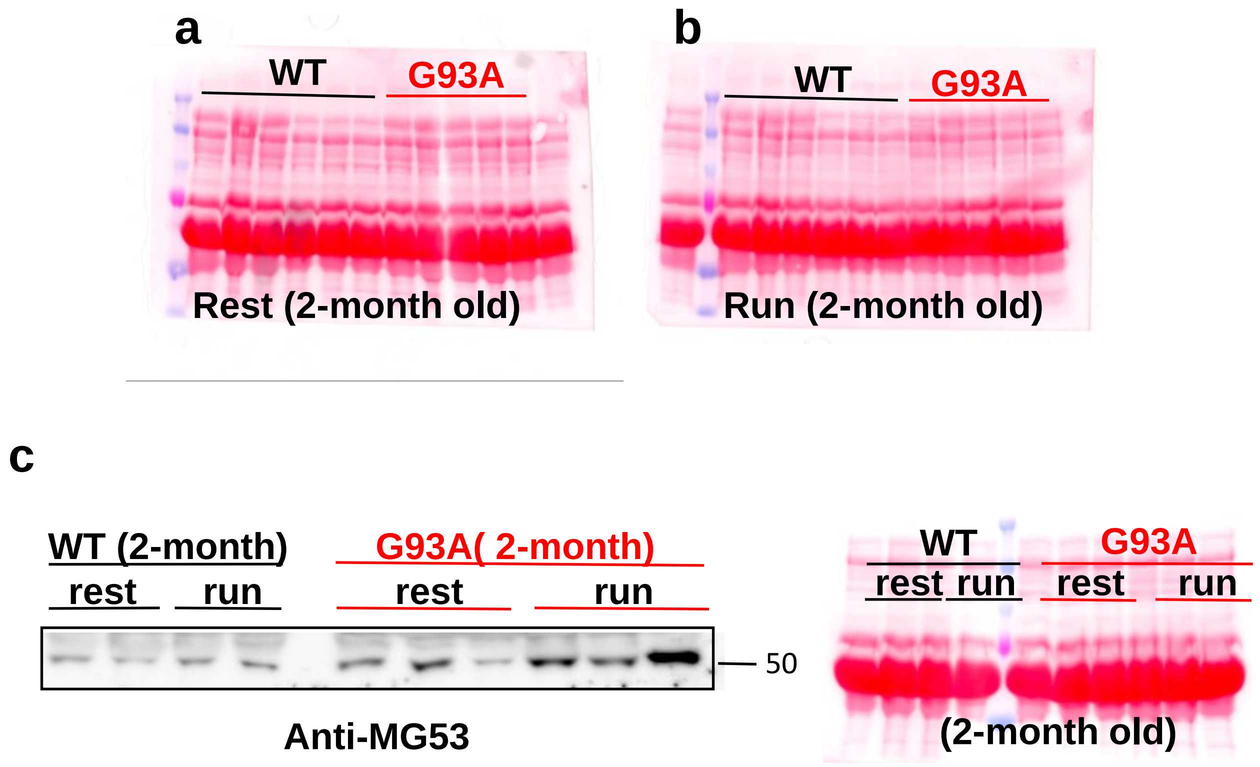

**Supplementary Fig. 2**(**a**,**b**) Ponceau S staining of the immunoblotting assay in **Fig. 1g** verifies equal loading amount of the serum samples. (**C**) Due to limited numbers of lanes in the acrylamide gel, the additional blotting assay of 2 WT and 3 G93A serum samples before and after running were evaluated in a separate gel. Data from this blot were also included in **Fig. 1g** quantification results.

#### Supplementary Fig. 3

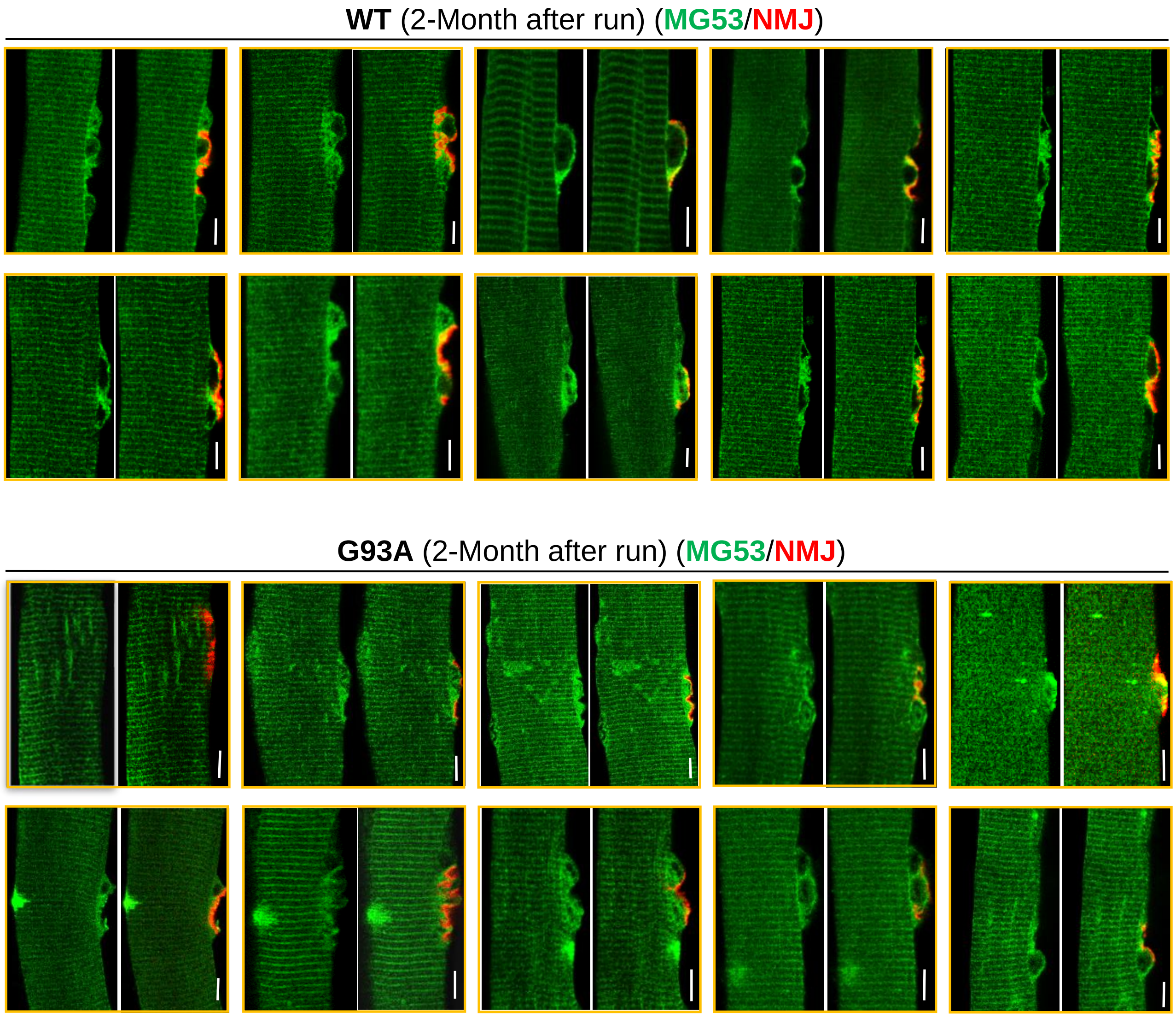

**Supplementary Fig. 3**. Additional representative images for **Fig. 2a**. FDB myofibers obtained from WT and G93A mice (2 months) subjected to 30 min running, were co-stained with MG53 antibody (green) and BTX (red). MG53 formed patches covering the NMJ site in both WT and G93A myofibers. However, intracellular MG53 aggregates started to appear near the site of NMJ in the G93A fibers. Scale bars: 20 µm.

#### Supplementary Fig. 4

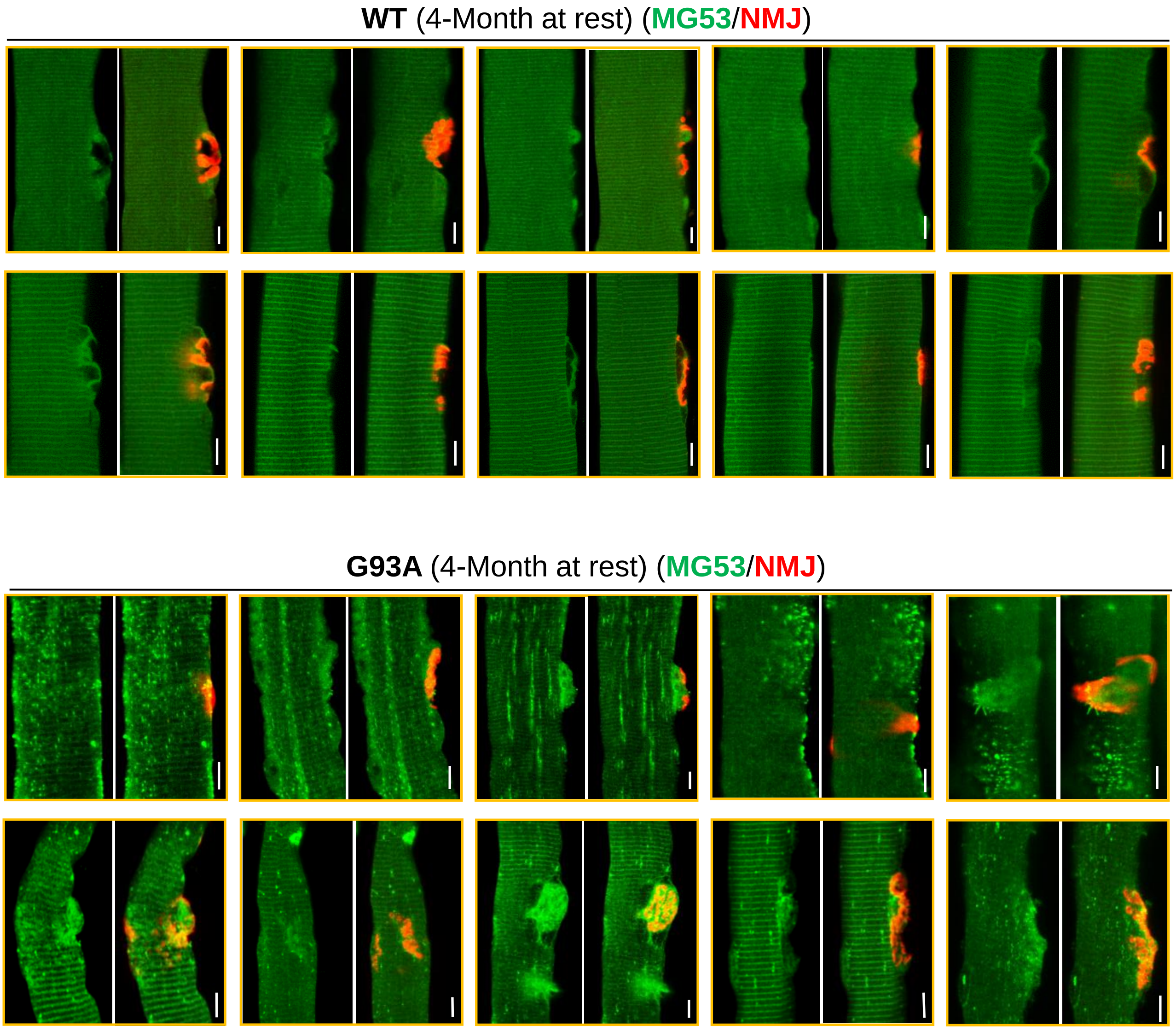

**Supplementary Fig. 4**. Additional representative images for **Fig. 2b**. FDB fibers derived from G93A mice at the advanced ALS stage (4-month old) at rest, were co-stained with MG53 antibody (green) and BTX (red). G93A fibers displayed extensive ectopic MG53 aggregates near the site of NMJ. Scale bars: 20 µm.

#### Supplementary Fig. 5

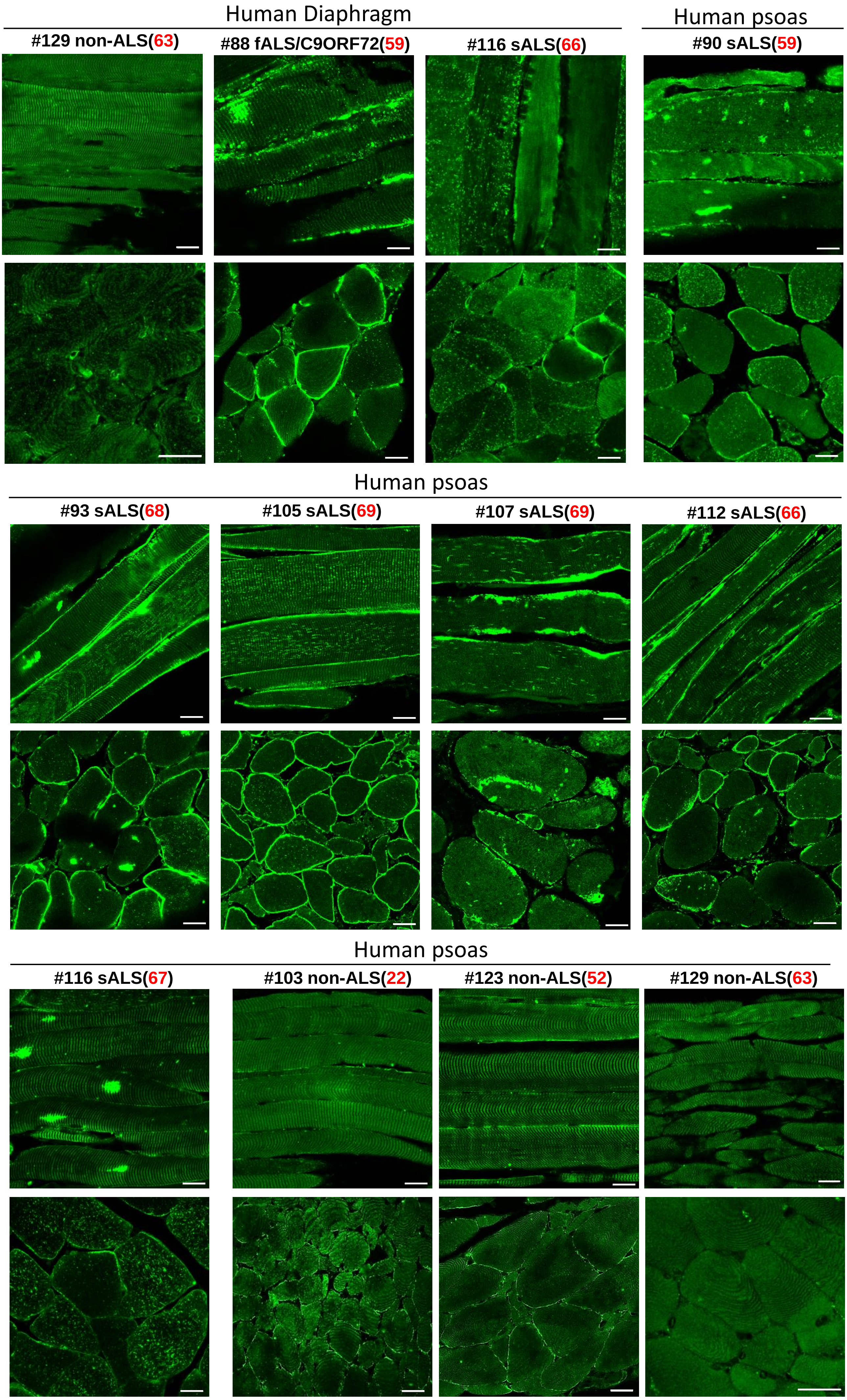

**Supplementary Fig. 5**. Additional images of anti-MG53 immunostaining of ALS and non-ALS human muscle samples. Bar: 20μm. The same as demonstrated in **Fig 3a**. both human ALS diaphragm and psoas muscles displayed extensive intracellular aggregates of MG53 or enhanced sarcolemma membrane localization. The age of patients was indicated with red fonts. Scale bars: 20 μm.

#### Supplementary Fig. 6

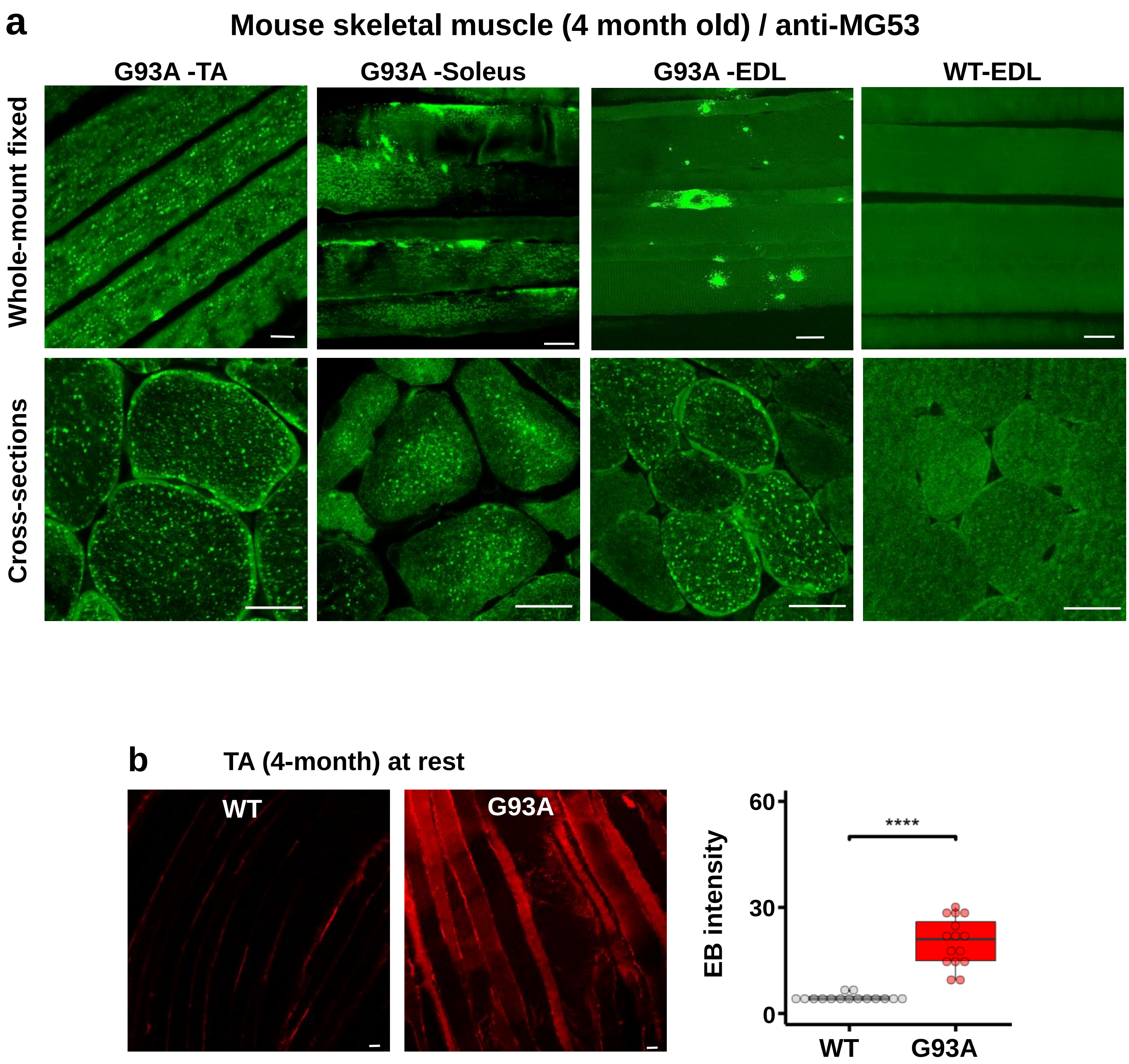

**Supplementary Fig. 6** (**a**) MG53 immunostaining of skeletal muscles isolated from multiple anatomic locations in 4-month old WT and G93A mice in cross sections and whole-mount preparations. There were prominent intracellular MG53 protein aggregates in all G93A muscles examined, while there was no apparent MG53 aggregation inside the WT muscles. (**b**) Representative images of TA muscle derived from G93A and WT mice (4-month old) at rest, which received EB injection 16 hours earlier. The EB intensity in TA muscle is significantly enhanced in G93A mice (WT: 4.46 ± 0.27, n = 15 from 2 mice *vs* G93A: 20.18 ± 1.77, n = 15 from 3 mice, ****P < 0.0001). Scale bars: 20 µm.

**Supplementary Table 1**: Demographics of human decedents

| ID | Race/ Gender | Age at onset | Age at death | Site of onset | Diagnosis | C9ORF72 expansion | SOD1 mutation |
|---|---|---|---|---|---|---|---|
| 38 | WF | 33 | 34 | Upper + lower extremities | fALS | Yes | No |
| 88 | WM | 57 | 59 | Bulbar | fALS + FTD | Yes | No |
| 90 | WF | 57 | 59 | Bulbar + Respiratory | sALS | No | No |
| 93 | WM | 65 | 68 | left hand | sALS | No | No |
| 102 | BF | 70 | 72 | Bulbar | sALS | No | No |
| 104 | WF | 69 | 71 | Bulbar | sALS | No | No |
| 105 | WM | 61 | 69 | left leg | sALS | No | No |
| 107 | WM | 67 | 69 | left hand | sALS | No | No |
| 112 | WM | 67 | 69 | right hand | sALS | No | No |
| 116 | WF | 64 | 67 | Left foot | sALS | No | No |

| ID | Race/ Gender | Age at death | Diagnosis | Cause of Death |
|---|---|---|---|---|
| 103 | WM | 22 | Non-Neurological Control | Metastatic synovial sarcoma (no CNS mets) |
| 123 | BM | 52 | Non-Neurological Control | Pancreatic Cancer (no CNS mets) |
| 129 | WF | 63 | Non-Neurological Control | Colon cancer with liver & lymph node mets (no CNS mets) |

**Supplementary Table 2**: Biospecimen information for human samples

| ID | Biospecimen type | Anatomic site | Vital state of patient | Collection | Type of stabilization | Type of long term preservation | Constitution of preservative | Storage duration | Shipping temperature | Composition Assessment & selection |
|---|---|---|---|---|---|---|---|---|---|---|
| 38 | Skeletal muscle | Diaphragm | Postmortem | Autopsy | Formalin fixed | Paraffin embedded | 10% neutral-buffered formalin | 21 years | 25°C | Clinical, genetic & neuropathological |
| 88 | Skeletal muscle | Diaphragm | Postmortem | Autopsy | Formalin fixed | Paraffin embedded | 10% neutral-buffered formalin | 1-7 years | 25°C | Clinical, genetic & neuropathological |
| 90 | Skeletal muscle | Psoas | Postmortem | Autopsy | Formalin fixed | Paraffin embedded | 10% neutral-buffered formalin | 1-7 years | 25°C | Clinical, genetic & neuropathological |
| 93 | Skeletal muscle | Psoas | Postmortem | Autopsy | Formalin fixed | Paraffin embedded | 10% neutral-buffered formalin | 1-7 years | 25°C | Clinical, genetic & neuropathological |
| 102 | Skeletal muscle | Diaphragm | Postmortem | Autopsy | Formalin fixed | Paraffin embedded | 10% neutral-buffered formalin | 1-7 years | 25°C | Clinical, genetic & neuropathological |
| 104 | Skeletal muscle | Diaphragm | Postmortem | Autopsy | Formalin fixed | Paraffin embedded | 10% neutral-buffered formalin | 1-7 years | 25°C | Clinical, genetic & neuropathological |
| 105 | Skeletal muscle | Psoas | Postmortem | Autopsy | Formalin fixed | Paraffin embedded | 10% neutral-buffered formalin | 1-7 years | 25°C | Clinical, genetic & neuropathological |
| 107 | Skeletal muscle | Psoas | Postmortem | Autopsy | Formalin fixed | Paraffin embedded | 10% neutral-buffered formalin | 1-7 years | 25°C | Clinical, genetic & neuropathological |
| 112 | Skeletal muscle | Psoas | Postmortem | Autopsy | Formalin fixed | Paraffin embedded | 10% neutral-buffered formalin | 1-7 years | 25°C | Clinical, genetic & neuropathological |
| 116 | Skeletal muscle | Diaphragm/ psoas | Postmortem | Autopsy | Formalin fixed | Paraffin embedded | 10% neutral-buffered formalin | 1-7 years | 25°C | Clinical, genetic & neuropathological |
| 103 | Skeletal muscle | Diaphragm/ psoas | Postmortem | Autopsy | Formalin fixed | Paraffin embedded | 10% neutral-buffered formalin | 1-7 years | 25°C | Clinical, genetic & neuropathological |
| 123 | Skeletal muscle | Psoas | Postmortem | Autopsy | Formalin fixed | Paraffin embedded | 10% neutral-buffered formalin | 1-7 years | 25°C | Clinical, genetic & neuropathological |
| 129 | Skeletal muscle | Psoas | Postmortem | Autopsy | Formalin fixed | Paraffin embedded | 10% neutral-buffered formalin | 1-7 years | 25°C | Clinical, genetic & neuropathological |

**Supplementary Table 3**: The survival days of individual tested G93A mice separated by gender

| Gender | Male | Male | Male | Male | Male | Male | | | T-test P | Chi square test P |
|---|---|---|---|---|---|---|---|---|---|---|
| Saline | 126 | 127 | 121 | 126 | 115 | | | | | |
| PEG-MG53 | 132 | 143 | 147 | 140 | 134 | 128 | | | <0.005 | |
| Between gender | | | | | | | | | | 1 |
| Gender | Female | Female | Female | Female | Female | Female | Female | Female | | |
| Saline | 137 | 122 | 131 | 119 | 128 | 119 | 119 | 124 | | |
| PEG-MG53 | 152 | 134 | 132 | 148 | 132 | 126 | 127 | | <0.05 | |

**Supplementary Table 3**: The survival days of individual tested G93A mice separated by gender. The Chi square test further confirmed that the significant difference between PEG-rhMG53 (PEG-MG53) and saline treated groups is independent of gender.

#### Supplementary Movie 1

Eight 3-month old G93A littermate mice were divided into two groups. Mice in the rhMG53 group (Right panel) received IV injection of PEG-rhMG53 (2 mg/kg) every other day for 30 days, while mice in Saline group (Left panel) received IV injection of saline every other day for 30 days. The videos were taken on the last day of the treatment.
